# Spontaneous alpha and theta oscillations are related to complementary aspects of cognitive control in younger and older adults

**DOI:** 10.1101/2020.04.09.033811

**Authors:** Grace M. Clements, Daniel C. Bowie, Mate Gyurkovics, Kathy A. Low, Monica Fabiani, Gabriele Gratton

## Abstract

The resting-state human EEG power spectrum is dominated by alpha (8-12 Hz) and theta (4-8Hz) oscillations, and also includes non-oscillatory broadband activity inversely related to frequency (1/*f* activity). Gratton (2018) proposed that alpha and theta oscillations are both related to cognitive control function, though in a complementary manner. Alpha activity is hypothesized to facilitate the *maintenance* of representations, such as task sets in preparation for *expected* task conditions. In contrast, theta activity would facilitate *changes* in representations, such as the *updating* of task sets in response to *unpredicted* task demands. Therefore, theta should be related to reactive control (which may prompt changes in task representations), while alpha may be more relevant to proactive control (which implies the maintenance of current task representations). Less is known about the possible relationship between 1/*f* activity and cognitive control, which was analyzed here in an exploratory fashion. To investigate these hypothesized relationships, we recorded eyes-open and eyes-closed resting-state EEG from younger and older adults and subsequently tested their performance on a cued flanker task, expected to elicit both proactive and reactive control processes. Results showed that alpha power and 1/*f* slope were smaller in older than younger adults, whereas theta power did not show age-related reductions. Resting alpha power and 1/*f* slope were predictive of proactive control processes, whereas theta power was related to reactive control as measured by the cued flanker task. All predictive associations were present over and above the effect of age, suggesting that these resting-state EEG correlates could be indicative of trait-like individual differences in cognitive control performance, which may be already evident in younger adults, and are still similarly present in healthy older adults.

## 1. Introduction

Three main features dominate the resting-state EEG power spectrum: alpha oscillations (8-12 Hz; e.g., Hanslmayr, Gross, Klimesch, & Shapiro, 2011; Jensen & Mazaheri, 2010; Mathewson et al., 2011), theta oscillations (4-8 Hz; e.g., Jaušovec, Jaušovec, & Gerlič, 2001; Pscherer et al., 2019;) and non-oscillatory broadband activity inversely related to frequency, known as 1*/f* slope (e.g., He, 2014). Other frequency bands have also been identified and investigated, such as beta, gamma, etc., but their amplitude is smaller. Alpha and theta oscillations have been extensively investigated in relation to cognition, whereas the relationship between 1*/f* activity and cognition is emerging (Cavanagh & Frank, 2014; Clayton, Yeung, & Kadosh, 2015; Cohen, 2014; Voytek et al., 2015). Alpha and theta power are most often measured during tasks to elucidate moment-to-moment neural variability yoked to certain stimuli or conditions. For instance, posterior alpha has been related to the inhibition of the processing of visual stimuli (e.g., Jensen & Mazaheri, 2010; Klimesch, Sauseng, & Hansilmayr, 2007; Lőrincz et al., 2009; Mathewson et al., 2009; 2011) and can be suppressed by incoming visual stimulation that needs attending (Morrell & Ross, 1953; Williamson et al., 1997). This suggests that alpha may be related to a processing mode geared at limiting the progression of perceptual information through the brain to avoid interfering with currently active representations.

In contrast, task-related activity in theta power shows marked, short-lived increases in response to stimuli with high levels of conflict or when task settings require updating (Cavanagh & Frank, 2014; Cavanagh et al., 2009; Clayton, Yeung, & Kadosh, 2015; Cohen, 2014; Cooper et al., 2016, Cohen & Donner, 2013). In this context, theta activity may be associated with the adjustment of settings related to how stimulus information needs to be processed (Cavanagh et al., 2009). Thus, in task-related conditions, both alpha and theta are thought to be associated with mechanisms regulating the flow of information, a set of processes often labeled cognitive control (for a review, see Gratton et al., 2018).

Providing a unified view of this evidence for alpha and theta, Gratton (2018) hypothesized that these oscillations exert complementary roles in cognitive control, with alpha associated with the maintenance of currently active representations (in order to protect their processing from interference; *proactive control,* Braver, 2012) and theta associated with the disruption/updating of representations, when attention needs to shift to incoming information (and alpha is therefore suppressed; *reactive control,* Braver, 2012). While largely proposed on the basis of stimulus-related activity, such complementary roles for alpha and theta could also occur spontaneously, outside of the influence of externally defined tasks, and therefore be related to trait-like *individual differences* in cognitive control. If this were the case, then alpha activity at rest could be *predictive* of the propensity of an individual to exert proactive control, which requires the maintenance of representations or goal-states. Similarly, the extent to which theta activity is exhibited at rest could be *predictive* of a person’s ability to detect and resolve interference, and therefore to exert reactive control. In addition, because cognitive control is known to vary with age (Braver & Barch, 2002, Braver, 2012; Bugg, 2014; Manard, Francois, Phillips, Salmon, & Collette 2017), we were also interested in determining whether aging would modulate these hypothesized relationships. Specific evidence for these hypotheses is reviewed in the remainder of this introduction.

### Alpha and theta power at rest

At rest, alpha power decreases with open eyes and is correlated with many cognitive processes, including working memory and IQ (Doppelmayr, Klimesch, Stadler, Pöllhuber, & Heine, 2002; Oswald et al., 2017). Further, it is well documented that alpha power decreases with age (e.g., Polich, 1997) and also in various stages of clinical impairments as dementia develops (Babiloni et al., 2006). Healthy older adults have been shown to have greater resting-state alpha power compared to those with Mild Cognitive Impairment (MCI), and they, in turn, have greater alpha power than those with Alzheimer’s Disease (AD; Moretti et al., 2004; Babiloni et al., 2006). Thus, resting-state alpha power has already been used as a biomarker for individual differences in older adults, due to its clinical relevance in distinguishing various degrees of age-related cognitive pathologies. Additionally, Mahjoory et al. (2019) have shown that resting-state alpha power in younger adults is related to attention span, indicating that alpha at rest may be a useful tool to understand cognitive variability in younger adults. By-and-large, these data indicate that high resting-state alpha power is associated with higher cognitive abilities, suggesting that resting-state alpha manifests a brain mechanism of significant importance for cognition.

The relationship between *resting-state* theta activity and cognitive control is less clear. Pscherer et al. (2019) found that individuals with low eyes-open resting-state theta power had poorly controlled conflict-related response inhibition during a Go/No-Go task compared to those with high resting theta power. They also found that participants with low resting-state theta power had more total task-based theta power on incompatible than compatible trials. This was not true for participants who had high resting-state theta power. These data suggest that resting theta power (just as theta power during tasks) is related to inhibitory control, particularly during conflict, and that theta at rest may predict cognitive control theta activity during a task. Surprisingly, however, theta at rest has also been negatively associated with lower IQ (Jaušovec, et al., 2001).

These seemingly contradictory findings suggest that, although theta oscillations may represent cognitive-control-related processes even in the absence of a task (but see Gordon et al., 2018), the exact nature of these processes is still unclear. They may reflect periodic disengagement from established representations to monitor the environment for changes, or the excessive occurrence of shifting or updating operations (Miyake & Friedman, 2012), which may characterize highly distractible individuals. Differently from alpha, the evidence for resting-state theta changes with age is mixed (Babiloni et al., 2006; Finnigan & Robertson, 2011). Theta power has been reported not to differ between healthy older adults and individuals with MCI or AD (Babiloni et al., 2006), but also to undergo a relative *increase* with disease progression (Kwak, 2006).

### 1/f activity

In recent years it has become clear that the resting-state EEG spectrum contains not only recurrent oscillatory activity, but also activity of a non-recurrent (or non-oscillatory) nature. These non-oscillatory phenomena typically produce a broadband effect, which is more visible at low than at high frequencies, likely because the longer the duration of these deflections, the greater their summation in the EEG spectrum. This type of activity is referred to as 1/*f* noise, 1/*f* slope or 1/*f* activity, because its power decreases as a function of frequency (*f*) following a power-law function (He, 2014; Voytek & Knight, 2015). Research on 1/*f* slope suggests that it could be related to task performance and cognitive state (Miller et al., 2015; Ouyang et al., 2020; Pertermann et al., 2019). It has also been recently shown that 1/*f* may be affected by age (Voytek et al., 2015; Dave et al., 2018;). Because 1/*f* is a substantial component of the resting EEG power spectrum, and may vary with age, we decided it would be important to separately estimate its role and predictive value in a set of exploratory analyses. Crucially, separating 1/*f* from the remainder of the spectrum is important to accurately estimate the power of oscillatory activity such as alpha and theta (Haegens, Cousijn, Wallis, Harrison, Nobre, 2014; Nikulin & Brismar, 2006; Voytek et al., 2015).

### Assessment of proactive and reactive control

To provide an independent assessment of an individual’s ability to exert proactive and reactive cognitive control, we employed a cued flanker task (Eriksen & Eriksen, 1974). In a flanker paradigm, a central target stimulus is flanked by irrelevant distractors that are either congruent (e.g., >>>>> or <<<<<) or incongruent with the target (e.g., <<><< or >><>>). The participant’s task is to ignore the flankers and respond based on the direction of the central arrow. In the *cued* version of this paradigm used in the current study, the reaction stimulus arrays are preceded by a cue that indicates the probability that the array will contain congruent flankers. The difference in performance between incongruent and congruent trials, known as the *congruency effect* (CE), reflects the extent to which distractor information is processed up to the point it influences responses. The less the distractor information is processed, the smaller the congruency effect, which is taken as a measure of *reactive control.* Smaller congruency effects indicate more effective reactive control.

Gratton, Coles, & Donchin (1992) showed that probability cues influence the size of the congruency effect, with cues indicating a high probability of congruency leading to larger congruency effects. In other words, when participants are warned that distractors are likely to be incongruent, they adopt a strategy that limits the influence of the distractor information, reducing the congruency effect. In contrast, when they are informed that irrelevant distractors are likely to be congruent, they adopt a strategy that allows for more distractor information to be processed, because on the majority of trials this information will facilitate performance. This is of course at the cost of hindering performance on the infrequent incongruent arrays, leading to a larger congruency effect in such instances. These effects are similar to the conflict adaptation effect (also called the congruency sequence effect, or the Gratton effect) which is characterized by a reduced congruency effect on trials immediately following an incongruent, as opposed to a congruent, array (e.g., Egner, 2007; Ullsperger, Bylsma, & Botvinick, 2005). We refer to the difference in probability cue-based congruency effects as the *conflict expectation effect* (CEE), which is taken here as a measure of *proactive control.* Age should lead to a reduction in the efficiency of both modes of control (i.e., an increase in CE and a decrease in CEE), although potentially less so for reactive control (Bugg, 2014). Using a cued-flanker task, we investigated the relationship between cognitive control processing and resting-state alpha power, theta power and 1*/f* slope in a sample of younger and older adults.

## 2. Materials and Methods

### 2.1 Participants

Twenty-one younger and 20 older adults were recruited and underwent the procedures described below^1^. Participants reported no history of psychiatric or neurological disorders and had no signs of dementia (scores ≥51 on the modified Mini-Mental Status Examination [mMMSE]; Mayeux, Stern, Rosen, & Leventhal, 1981), or depression (younger adults assessed with the Beck’s Depression Inventory, Beck, Steer, & Brown, 1996; and older adults with the Geriatric Depression Scale, Yesavage et al., 1983; Yesavage & Sheikh, 1986). The study received ethical approval from the Institutional Review Board at the University of Illinois Urbana-Champaign. All participants provided written informed consent and were compensated for their time.

One younger adult and one older adult were excluded for excessive EEG artifacts: eye movements, muscle activity, and/or amplifier saturation. The remaining 20 younger adults (age range = 18-30, 14 females) and 19 older adults (age range = 65-80, 11 females) constituted the final sample. See Table 1 for age-group characteristics. Older and younger adults were well matched in cognitive status, although older adults, as is typical, had a slight advantage over younger adults in tests relying on vocabulary knowledge^2^, *t*(35) = 4.29, *p* = .0001. Older adults had slightly more years of education, *t*(37) = −2.15, *p* = .04, as expected, given that most younger adults were college students. Older adults also had higher composite age-adjusted IQ, *t*(37) = −2.25, *p* = .03, than younger adults.

**Table 1.** Descriptive characteristics of the sample. Mean (SD).

|  | **Younger Adults** | **Older Adults** | ***p*-value** |
| --- | --- | --- | --- |
| N | 20 (14 females) | 19 (11 females) |  |
| Age (years) | 22.4 (3.3) | 69.4 (4.3) | < .001 |
| Education (years) | 15.6 (2.4) | 17.3 (2.4) | .038 |
| IQ (age-adjusted)* | 115.2 (10.7) | 124.0 (15.4) | .030 |
| Shipley's Vocabulary Scale | 31.4 (3.4) | 36.1 (3.2) | < .001 |

*\*Kaufman Brief Intelligence Test – 2<sup>nd</sup> edition*

### 2.2 Data Acquisition and Analysis

Participants underwent two separate sessions: a resting-state EEG recording session, followed by a behavioral testing session.

#### 2.2.1 Resting-State EEG

Participants sat in a dimly lit, sound- and electrically attenuated recording chamber, and were instructed to sit quietly and not think about anything in particular. Each session included a 1-minute recording of resting EEG with eyes-open followed by 1-minute with eyes-closed. These recording periods were conducted at the beginning of a recording session that involved other experiments that will not be reported here.

#### 2.2.2 EEG Recording and Analysis

EEG and EOG were recorded continuously from 64 active electrodes in an elastic cap (Acti-Cap) using BrainAmp amplifiers (BrainVision Products GmbH). EEG was recorded from scalp electrodes referenced to the left mastoid, with off-line re-referencing to the average of the two mastoids. Two electrodes placed above and below the left eye measured vertical EOG to detect blinks and vertical eye movements. Two electrodes placed approximately 1 cm to the left and right of the outer canthi of the eyes measured horizontal eye-movements (saccades). Impedance was kept below 10kΩ. The EEG was filtered on-line using a 0.1-250 Hz bandpass and sampled at 500Hz.

Off-line EEG processing was done using EEGLAB Toolbox (version: 13.6.5b; Delorme & Makeig, 2004), ERPLAB Toolbox (version: 6.1.3) and custom Matlab16a scripts (The MathWorks, Inc., Natick, Ma, USA). A 30 Hz low-pass filter was applied. The data were epoched into 4096 ms contiguous segments to facilitate usage of our artifact detection scripts. Epochs with amplifier saturation were discarded. Ocular artifacts were corrected using the procedure described in Gratton et al. (1983). After eye movement correction, epochs with voltage fluctuations exceeding 200μV were excluded from further analysis to minimize the influence of any remaining artifactual activity. This resulted in the exclusion of 1.26% of the epochs.

Power spectral densities were determined using a fast Fourier transform with Welch’s method at parietal (Pz, POz, P1, P2, PO3, PO4) and frontocentral (Fz, FCz, Cz, CPz, FC1, FC2, C1, and C2) electrode clusters. The segments were zero-padded and multiplied with a Hamming taper, with 0% overlap. These clusters were selected because alpha power is largest at parietal locations (e.g., Haegens, Cousijn, Wallis, Harrison, & Nobre, 2014), and 1/*f* slope and theta are typically largest at frontocentral locations (Pertermann, Mückschel, Adelhöfer, Ziemssen, & Beste, 2019). This was confirmed in our sample by a review of topographical plots.

For the purposes of the current study, we modeled the observed power spectrum as formed by the sum of non-oscillatory activity (1/*f*), alpha and theta oscillatory peaks, and noise using the following equation. Because there was no observable peak in the beta band, beta was not included in the model:

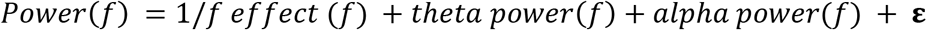

where *f* is frequency and **ε** is noise.

To ensure that our measures of alpha and theta oscillatory power were not confounded with non-oscillatory activity, the 1*/f* component was removed from the spectrum prior to calculating mean narrowband power. We first modeled the 1*/f* component on the raw power spectrum, excluding 4-13 Hz, which represents most oscillatory power in the spectrum, using a least-squares linear regression with 1/*f* predicting power (see Figure 1; frequencies up to the Nyquist frequencies were used to model *1/f,* frequencies up to 30 Hz were plotted). The slope of the 1*/f* component – i.e., the ß weight of predictor 1/*f* in the equation – was retained for analysis. Note that the exponent of the *f* predictor was fixed at −1 (i.e., *f*^−1^). In other words, we did not use a log-log transform to fit the exponent of the 1/*f* function to the data, as done in other studies (e.g., Voytek et al., 2015). Rather, we transformed the frequency values into their inverse, and then regressed the power values for each frequency onto this new axis (excluding power values between 4 and 13 Hz). This procedure was used because it is less sensitive to very small variations in power for high frequencies, which in the EEG spectrum (compared to the ECoG spectrum) have very low power. Because of these procedures, the interpretation of the 1/*f* slope as computed in this fashion is different than that used by Voytek and colleagues (2015). Rather than reflecting the shape (i.e., the exponent) of the 1/*f* function, the slope of the 1/*f* function in the current study is related to *power at very low frequencies* - and thus corresponds more closely to the *intercept* of the 1*/f* function in log-log space as reported by Voytek and coll. (2015).

The 1*/f* trend was then subtracted from the spectrum, allowing for more reliable alpha and theta estimates (Nikulin & Brismar, 2006; Haegens et al., 2014). Mean alpha and theta power were quantified on the detrended spectra and then log-transformed. The log transformed data were used for analysis because power has a positively skewed distribution. Using the logarithm attenuates the skewness of the power distributions and allows for an ANOVA to be performed without violating the assumption of normality. Similarly, the 1*/f* slope was also log-transformed.

**Figure 1:**
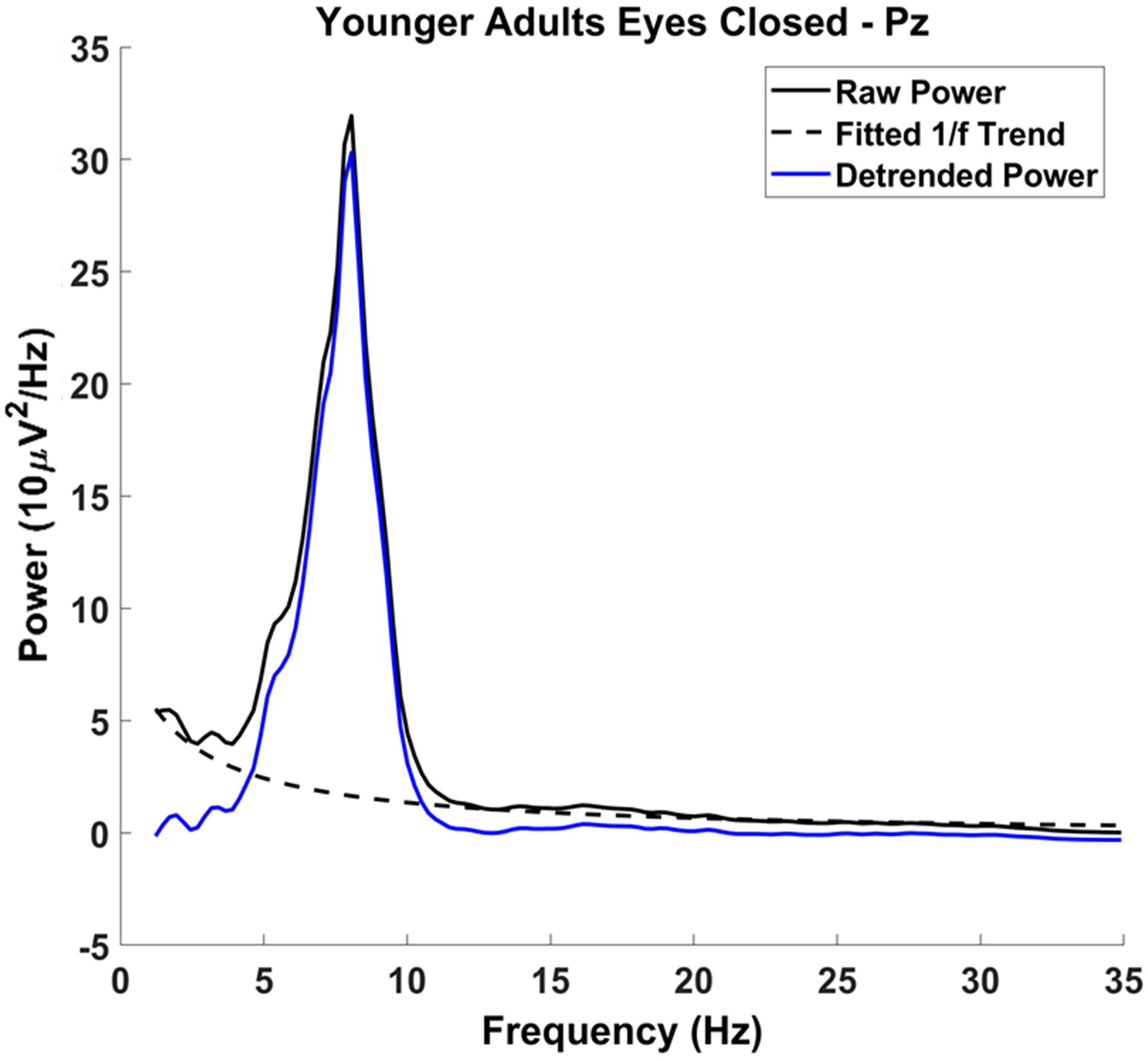
Detrending Procedure: Raw power spectrum (solid black line), with the characteristic 1/*f* phenomenon across the spectrum. The 1/*f* phenomenon was modeled excluding the frequencies 413 Hz (dashed black line) and was subtracted from the raw power spectrum, resulting in the detrended power spectrum (solid blue line). Alpha and theta measurements were made on the detrended power spectrum.

Assessment of alpha power scalp topographies separately at 8, 9, 10, 11, and 12 Hz revealed low power across the scalp at 8 Hz and 12 Hz in both younger and older adults. Thus, we estimated alpha power based on the average power between 9-11 Hz. For theta, which did not show a prominent peak in the spectra, we used a broader frequency window encompassing the full, canonical theta band, 4-8 Hz.

#### 2.2.3 Behavioral Task Session, Stimuli, and Related Analyses

At the beginning of testing, participants were seated 100 cm in front of a computer monitor centered at eye-level and were read instructions by the experimenter to supplement on-screen instructions. The imperative stimulus consisted of five horizontal arrows that were either congruent (<<<<< or >>>>>) or incongruent (<<><< or >><>>) on any given trial. Participants indicated, as quickly and accurately as possible, which direction (left or right) the central, target stimulus was pointing by pressing one of two keypads located on either side of the participant. Stimulus-response mapping was constant across all participants (i.e., a left-pointing target stimulus always required a left-button press, and vice versa) to eliminate the possibility of a confounding Simon effect in some participants (Simon, 1969).

Three neutral, low-arousing images of inanimate objects (fire hydrant, dresser, and screw, from the International Affective Picture System database, IAPS; Lang, Bradley, & Cuthbert, 2008) served as cues and preceded the imperative stimulus array. These images represented a 75% (predict-congruent; PC), a 50% (predict-neutral; PN), and a 25% (predict-incongruent; PI) probability of a congruent stimulus array, respectively. The three cue types were equiprobable and participants were explicitly told the congruency probability represented by each cue prior to commencing the task. PC and PI cue images were counterbalanced across subjects.

Each trial began with a 499 ms cue, followed by a 999 ms fixation. Then, the imperative stimulus appeared for 149 ms and was followed by 1848 ms of fixation before the onset of the next trial. The response window began with the onset of the imperative stimulus and continued until the onset of the next cue (i.e., the next trial). The global probability of a congruent trial within each block was 50%. The imperative stimulus arrays were presented in white typeface on a black computer screen and subtended 2.23° × 0.46°. Each cue overlaid a gray background with uniform dimensions such that each composite image subtended 6.98° × 5.35°. All stimuli were presented on a monitor (19-in. CRT, refresh rate 60 Hz, screen resolution 1280 × 960; Dell Computer, Round Rock TX) using the E-Prime 2.0 software (Psychology Software Tools, Pittsburgh, PA).

Accuracy feedback was displayed on-screen at the end of each block. If accuracy was below 75% across all trial types, participants saw a message that read “respond more slowly and more accurately.” If they scored between 75 and 95%, they saw “continue to respond as quickly and accurately as you can.” If they scored above 95%, they saw “respond more quickly”. The feedback was designed to encourage participants to prioritize speeded responses and elicit a reasonable number of errors, a requirement for accurately assessing speed of processing. Participants could take breaks between blocks, as needed.

Before the flanker task, younger adults completed 96 practice trials at the experimental speed. Older adults completed two sets of practice trials. Additional practice was added for older adults to offset difficulties (apparent in preliminary data) for them to complete the task at the experimental speed. As such, we added a slower-paced practice block to familiarize this group with the task. In the first set (48 trials), the inter-stimulus interval (ISI) was increased by 30%, but the cue and imperative stimulus presentation times remained at experimental speed. In the second set (96 trials), each trial ran at the experimental speed. Subsequently, all participants completed three blocks of 288 experimental trials.

Incorrect trials and all trials with reaction times ≤ 200 ms (i.e., fast guesses) were discarded before statistical analysis. Data were collapsed across target stimulus direction (i.e., response hand) to create six trial-types: 3 cue types (PC, PI, PN) × 2 flanker congruency conditions (congruent, incongruent). In addition to recording reaction time (RT) and calculating error rates for each trial-type, the inverse efficiency score (IES), an integrated measure of RT and accuracy, was also calculated (Townsend & Ashby, 1978, 1983; Bruyer & Brysbaert, 2011). IES is insensitive to speed-accuracy tradeoffs and results in a measure of RT that is not biased by fast decisions. IES is computed by dividing the mean RT of correct responses by the proportion of correct responses for each trial type (IES = RT/proportion correct), thereby providing an index of processing speed that estimates the “true” processing speed when the effects of speed-accuracy tradeoffs are minimized.

For our measure of reactive control, we calculated the congruency effect (CE) by subtracting congruent trials from incongruent trials (Incongruent - Congruent), so smaller differences in RT, error rate, and IES indicate greater reactive control. For the measure of proactive control, we calculated the conflict expectation effect (CEE) by subtracting the predict incongruent CE from the predict congruent CE (PC_CE_ - PI_CE_). Here, a larger difference indicates greater proactive control.

### 2.4 Statistical Analyses

Two-way mixed ANOVAs were computed, with age (young, old) as a between-subjects factor, eye status (open, closed) as a repeated measure, and alpha power, theta power, or 1*/f* slope as dependent variables. Alpha analyses were restricted to the parietal cluster, whereas theta and 1*/f* analyses were restricted to the frontal cluster. Spearman’s *rank-order* correlation coefficients were computed between alpha power, theta power, 1*/f* slope and behavioral performance measurements (CE and CEE for error rate, RT, and IES) across participants, in order to assess the effects of individual variability in restingstate EEG parameters and subsequent cognitive control processing. Spearman’s rho was used instead of Pearson’s *r* because assessment of normality with the Shapiro-Wilk test indicated that the behavioral measures were not normally distributed. These correlations were calculated both with and without partialing out the effects of age. The CEs were assessed with age (young, old), cue-type (PC, PI, PN), and trial-type (congruent, incongruent) in three-way mixed ANOVAs. The CEE was assessed by comparing the predict-incompatible to the predict-compatible dependent variables (error rate, RT, and IES) in a 2 x 2 ANOVA with age as the between-subjects factor. To address the non-normality, behavioral data were log-transformed prior to computing the ANOVAs. Significance levels were corrected for multiple comparisons as noted in the results.

Given the relatively small sample size, we computed intrasubject reliability for alpha and theta power and 1*/f* slope across the recording period, to show the consistency of the measurements. This was done by comparing alpha power and theta power (after removing the 1/*f* effect) and 1/f slope during even and odd epochs (each 4096 ms in length). Spearman’s rho was calculated between these values for even and odd epochs for each participant.

## 3. Results

### 3.1 EEG Power Spectrum Decomposition

After decomposing the power spectrum into alpha power, theta power and 1*/f* slope we quantified the variance in the power spectrum captured by these components across participants. To do this, we partitioned each individual’s power spectrum into four components: the power accounted for using the 1/*f* slope, alpha power, theta power, and the remaining, residual power not captured by the model. Together, these should capture 100% of the variance in the power spectrum, given that power itself is a measure of variance of the EEG time-series. Taking the sum of power accounted for with 1*/f* slope, alpha, and theta and dividing it by the sum of all four components will yield the amount of variability in the power spectrum accounted for by the non-oscillatory and main oscillatory features of the EEG power spectrum. This was done for each participant and then averaged across participants to give an overall estimate of captured variance. With eyes open, these three parameters (alpha power, theta power, and 1*/f* slope) accounted for 90% of the spectral variance at the parietal electrode cluster and 93% at the frontocentral cluster. Alpha, theta, and 1*/f* slope accounted for slightly less variance with eyes closed: 89% at both the parietal and frontocentral electrode clusters. Therefore, although the EEG signal is rich and complex, it can be largely described by these parameters. Power spectra (after removing 1*/f* slope) for each electrode cluster are shown in Figure 2.

Alpha, theta and 1*/f* scalp distributions are presented in Figure 3. As shown in these maps, younger and older adults had a posterior alpha scalp distribution, and a more anterior theta, as typically observed. The scalp distribution of 1/*f* had both anterior and posterior aspects. All effects were larger around the midline.

**Figure 2:**
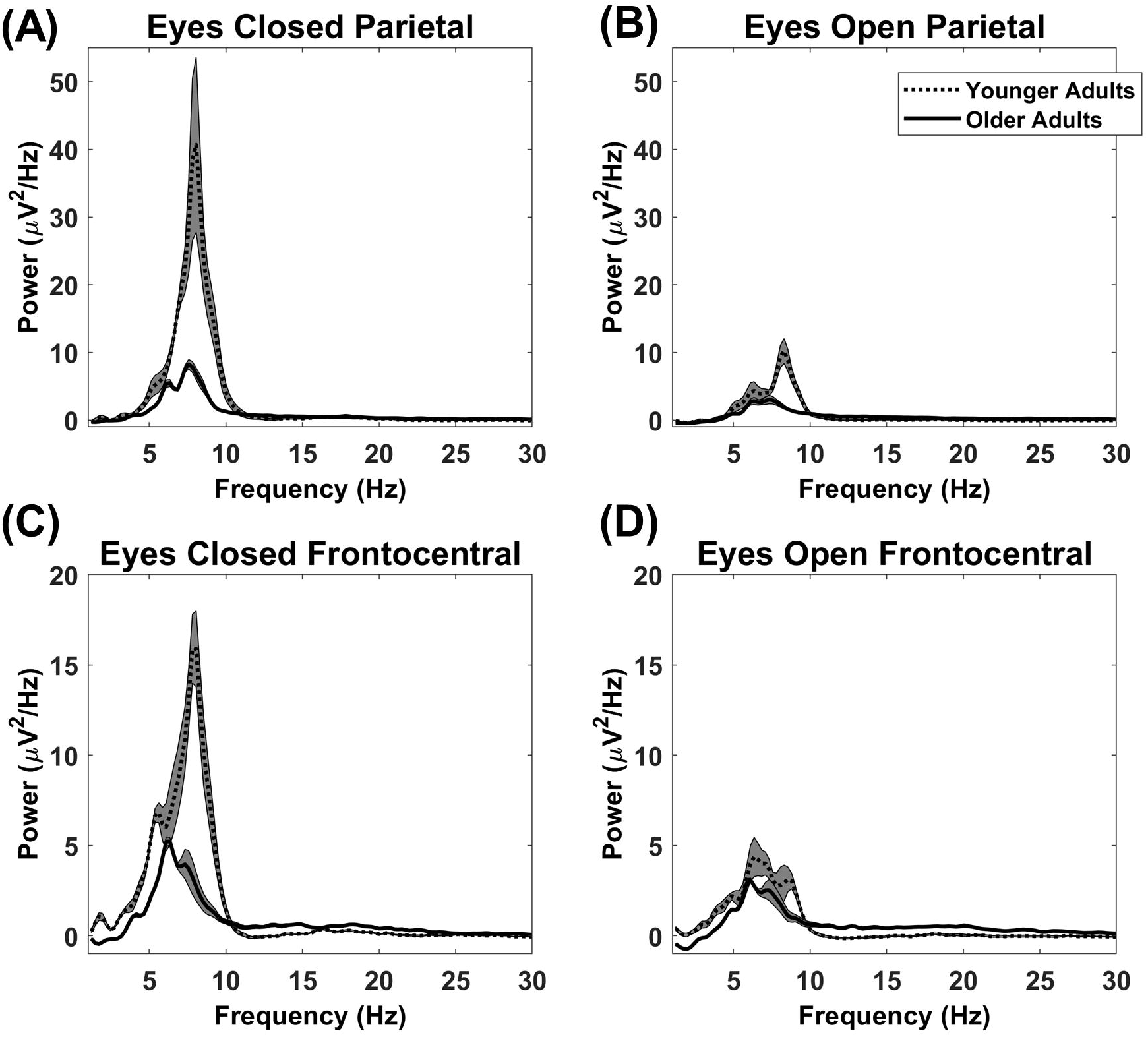
Detrended Power Spectra: Detrended power spectra at the parietal (A-B) and frontocentral (C-D) electrode clusters with eyes closed (A, C) and eyes open (B, D). Shaded gray areas indicate +/- the standard error. Note that younger adults had greater alpha power than older adults with both eyes open and eyes closed. However, theta power was not significantly different between younger and older adults. Both alpha and theta powers were reduced by opening the eyes.

**Figure 3:**
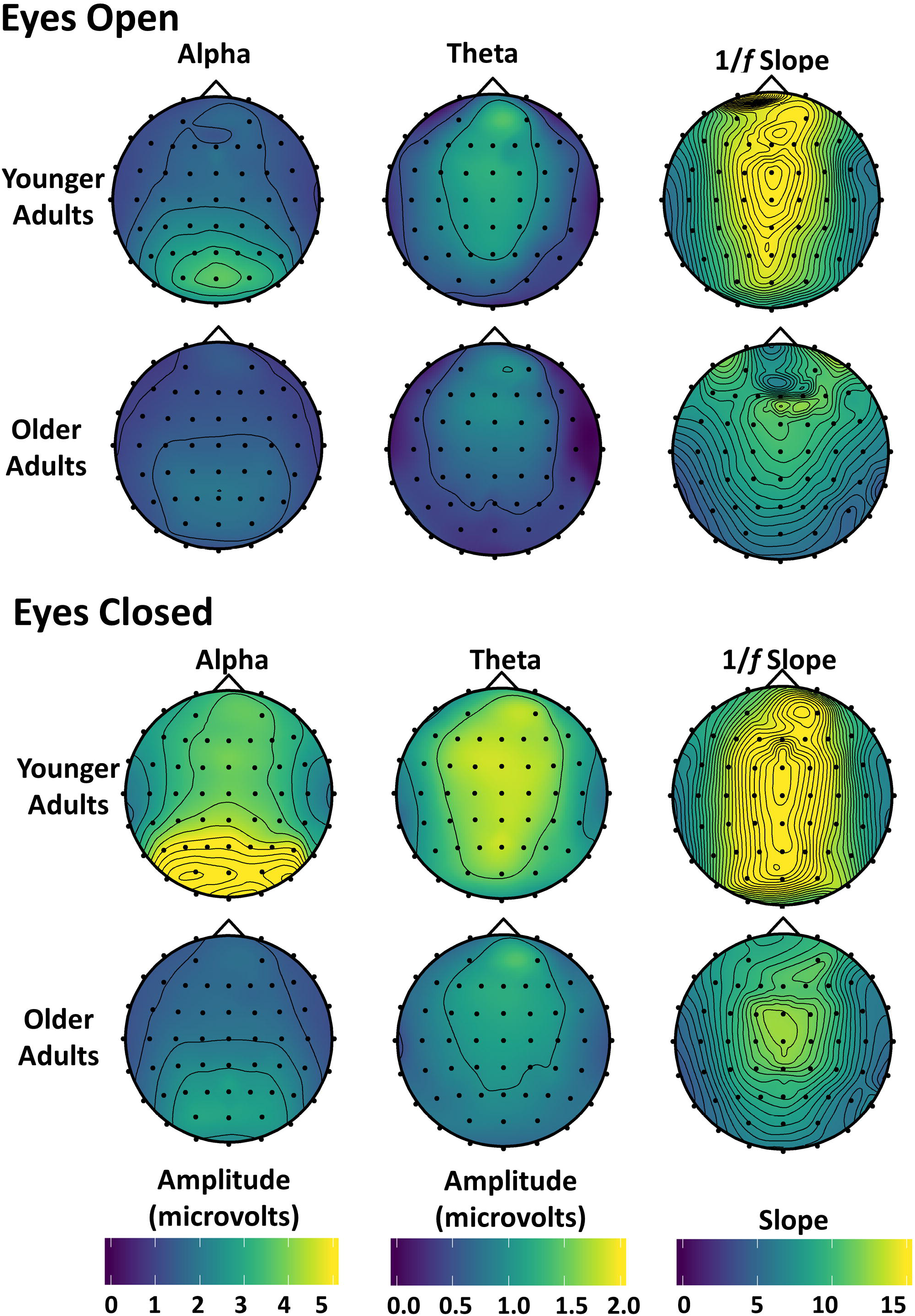
Topographic maps of alpha, theta, and 1*/f* slope: Alpha and theta amplitude, and 1/*f* slope with eyes open (top) and closed (bottom) are shown for both younger and older adults. Amplitude was used instead of power because power differences between younger and older adults were too large to be plotted meaningfully using the same scale.

Alpha power at the parietal cluster was positively correlated with theta power at the frontocentral cluster with eyes open, *r*(37) = .344, *p* = .032, [*r*(36) = .345, *p* =.034 after partialing out age], and with eyes closed, *r*(37) = .423, *p* = .007, [*r*(36) = .386, *p* = .017 after partialing out age]. Thus, theta and alpha power were correlated with each other at rest, both before and after accounting for age effects and in both eye conditions.

Neither alpha nor theta power were correlated with 1*/f* slope [alpha, eyes open, *r*(37) = .087, *p* = .600 or eyes closed, *r*(37) = .172, *p* = .295; theta, eyes open, *r*(37) = .185, *p* = .260, or eyes closed, *r*(37) = .132, *p* = .423]. All of these correlations remained non-significant after partialing out age.

### 3.2 Reliability Analyses Within Individuals

To show the consistency of the measurements, we computed intrasubject reliability for alpha and theta power and 1/*f* slope across the recording period, by comparing even and odd epochs (each 4096ms in length).^3^ We found that alpha power was highly consistent within individuals both with open (*r_s_* (37) = .922, *p* < .00001, average rho across participants) and closed eyes (*r_s_* (37) = .980, *p* < .00001, average rho across participants). Similarly, theta power was highly reliable both with open (*r_s_*(37) = .774, *p* < .00001) and closed eyes (*r_s_* (37) = .816, *p* < .00001). Lastly, 1/*f* slope was also consistent within individuals with open (*r_s_* (37) = .931, *p* < .00001) and closed eyes (*r_s_* (37) = .854, *p* < .00001).

### 3.3 Effects of age and eye status on alpha, theta, and 1/f

A 2(age) × 2(eye status) mixed effects ANOVA was run for alpha power at the parietal electrode clusters. As expected, alpha power was greater with closed compared to open eyes, *F*(1, 37) = 24.865, *p* < .001, partial η^2^ = .402 (see Figures 2-3). Although there was no main effect of age *F*(1, 37) = 1.056, *p* = .311, partial η^2^ = .208, there was an age by eye status interaction, *F*(1, 37) = 10.260, *p* = .003, partial η^2^ = .217. Assessing the simple effects with eyes open, there was little difference in alpha power between younger and older adults, *F*(1, 37) = .255, *p* = .616, partial η^2^ = .007 but when they closed their eyes, younger adults had significantly higher alpha power than older adults, *F*(1, 37) = 11.225, *p* = .002, partial η^2^ = .233. This may indicate that older adults modulate their resting-state alpha power less than younger adults, which, in turn, may reflect a weakening of the neural system giving rise to the alpha rhythm, or of the mechanisms for controlling it.

A similar analysis was run for theta power measured at the frontocentral cluster. The results indicated a main effect of eye status, *F*(1,37) = 25.438, *p* < .001, partial η^2^ = .407 with greater power with closed than open eyes. However, there was no significant difference between the two age groups, *F*(1,37) = 0.064, *p* = .801, partial η^2^ = .002, and the interaction between age and eye condition was not significant, *F*(1,37) = 2.850, *p* = .100, partial η^2^ = .072. Thus, the data suggest no effect of age on theta power at rest.

The 1*/f* slope was analyzed with a 2(age) × 2(eye status) mixed ANOVA at the frontocentral cluster to determine potential group differences and the effect of eye condition on 1/*f* slope. The ANOVA revealed that 1*/f* slope was greater with closed than open eyes, *F*(1, 37) = 8.223, *p* = .007, partial η^2^ = .182. There was also a main effect of age, *F*(1, 37) = 12.363, *p* = .001, partial η^2^ = .250, such that younger adults had higher levels of 1*/f* activity than older adults and this effect did not interact with eye status, F(1, 37) = .173, p = .680. An age-related reduction in 1*/f* slope had been previously shown during visual working memory tasks (Voytek et al., 2015; note however, that in their study this was labeled 1/*f* intercept, see Methods section) and language processing tasks (Dave et al., 2018), but to our knowledge this is the first time this age-related reduction has been demonstrated at the scalp in the absence of an explicit cognitive task.

We performed some additional ancillary analyses, reported in **Supplementary Materials**, to establish whether the timing patterns of theta and alpha occurrence at rest are similar to those observed during tasks. Specifically, we sought to investigate whether frontocentral theta *at rest* occurs in short bursts, as it typically does in tasks in response to the onset of attention-catching stimuli (Cavanagh & Frank, 2014; Cohen & Donner, 2013). Task-recorded alpha, instead, is often present before stimulus onset, with theta bursts typically occurring simultaneously with the alpha suppression that follows stimuli that capture attention. Thus, we predicted alpha to be present in a higher proportion of the time during the resting-state recording than theta. The results of these supplementary analyses are overall consistent with these hypotheses, supporting the idea that similar temporal patterns characterize the resting-state spectrum for theta and alpha compared to their task-based counterparts.

### 3.4. Behavioral Effects

Evidence from behavioral ANOVAs replicated the well-established CE for error rate [*F*(1, 37) = 24.708,*p* < .001, partial η^2^ = .381], RT [*F*(1, 37) = 272.342,*p* < .001, partial η^2^ = .88], and IES [*F*(1, 37) = 147.599, *p* < .001, partial η^2^ = .800]. There was a main effect of age on the CE when measured with RT, *F*(1, 37) = 22.514, *p* < .001, partial η^2^ = .378 and IES *F*(1, 37) = 22.797, *p* < .001, partial η^2^ = .381 such that older adults had a larger CE than younger adults.

The probability cues also influenced the size of the CE. This was reflected in a CEE for RT [*F*(1, 37) = 6.042, *p* =.035, partial η^2^ = .140] and IES [*F*(1, 37) =5.070, *p* = .047, partial η^2^ = .121], where CE’s were larger following predict congruent compared to predict incongruent trials. The CEE was not apparent for error rate, *F*(1, 37) = 2.412, *p* = .177, partial η^2^ = .061 and there was no age by CEE interaction for any of the 3 behavioral measures: error rate [*F*(1, 37) < 0.001, *p*> .999, partial η^2^ = 0.0], RT [*F*(1, 37) = 1.853, *p* = .222, partial η^2^ = .048] or IES [*F*(1, 37) = 0.001, *p*> .999, partial η^2^ = 0.0]. All reported *p*-values for the predicted main effects in this section were corrected for multiple comparisons using FDR.

### 3.5 Correlations between Cognitive Control and Resting-State Alpha Power, Theta Power and 1/f Slope

Critically, the current study explored the relationship between alpha power, theta power and 1*/f* slope and proactive and reactive cognitive control processes. The results reported here are exclusively for the eyes open condition, as the eyes closed condition did not predict behavior (it should be noted that this is not surprising, as the behavioral measures were obtained with the eyes-open). All correlations were corrected for multiple comparisons using family-wise FDR to control the expected proportion of false positives in our results. As shown in Figure 4A, alpha power predicted the size of the CEE for IES, *r_s_* (37) = .491, *p* = .024. This is consistent with the prediction that alpha is related to individual differences in proactive control. Alpha predicted RT CEE prior to multiple comparisons correction, *r_s_* (37) = .321, *p* = .046, but not after: *p* = .183. The relationship between IES CEE and alpha remained significant even after partialing out the effects of age, IES CEE: *r_s_* (36) = .487, *p* = .024, suggesting that the ability to engage proactive control is not driven by age per se, and may instead reflect individual differences in proactive cognitive control capacity that may be maintained in healthy aging. In contrast, alpha power did not predict the size of the IES CE, *r_s_* (37) = .118, *p* = .630 (with or without partialing out the effects of age). Thus, alpha power appears specifically related to proactive control abilities rather than to cognitive control in general, with greater proactive control in individuals with larger alpha power at rest.

For resting theta power, there was a marginal positive relationship with the error rate CE, *r_s_*(37) = .406, *p* = .06, which varied little when age was partialed out, *r*(36) = .410, *p* = .06. This effect was not present for RT CE, *r_s_* (37) = .020, *p* = .902 or with IES CE, *r_s_* (37) = .306, *p* = .183. Theta power was not significantly correlated with the CEE for any of the behavioral measures: error rate, *r_s_*(37) = .226, *p* = .332, RT *r_s_* (37) = −.040, *p* = .883, or IES, *r_s_* (37) = .182, *p* = .434, and remained nonsignificant after partialing out age. These results indicate a modest relationship between theta activity at rest and reactive, but not proactive, control. Interestingly, the data indicate that individuals with high theta power at rest are more affected by distractors than individuals with low theta power. These data are shown in Figure 4B.

It is important to investigate the dissociation between both alpha and theta power and their relationships to proactive and reactive control, respectively. Therefore, partial correlation analyses were conducted to further illustrate the effect of alpha power on proactive control processes while holding theta power constant and the effect of theta power on reactive control processes while holding alpha power constant. As throughout this section, these analyses were conducted on the eyes-open measurements and corrected for multiple comparisons using family-wise FDR. Alpha power was positively related to the IES CEE [*r_s_* (36) = .464, *p* = .018] and the RT CEE [*r_s_* (36) = .368, *p* = .046] after partialing out the effects of theta power. As before, the error rate CEE was not related to alpha power, even after partialing out theta power [*r_s_* (36) = .198, *p* = .281]. The relationship between theta power and reactive control processes also varied little after partialing out the effects of alpha power. As before, there was a positive relationship between theta power and the error rate CE [*r_s_* (36) = .402, *p* = .036,]. Taken together, these results indicate that alpha power is indeed related to proactive control processes, in a way that is independent from the contributions of theta power, and, conversely, that theta power may be related to reactive control processes, in a way that is independent from the contributions of alpha (although this result is only marginally significant). This suggests that alpha and theta power *at rest* are uniquely related to proactive and reactive control, respectively. As such, they could be used as separable biomarkers for these two aspects of cognitive control.

**Figure 4:**
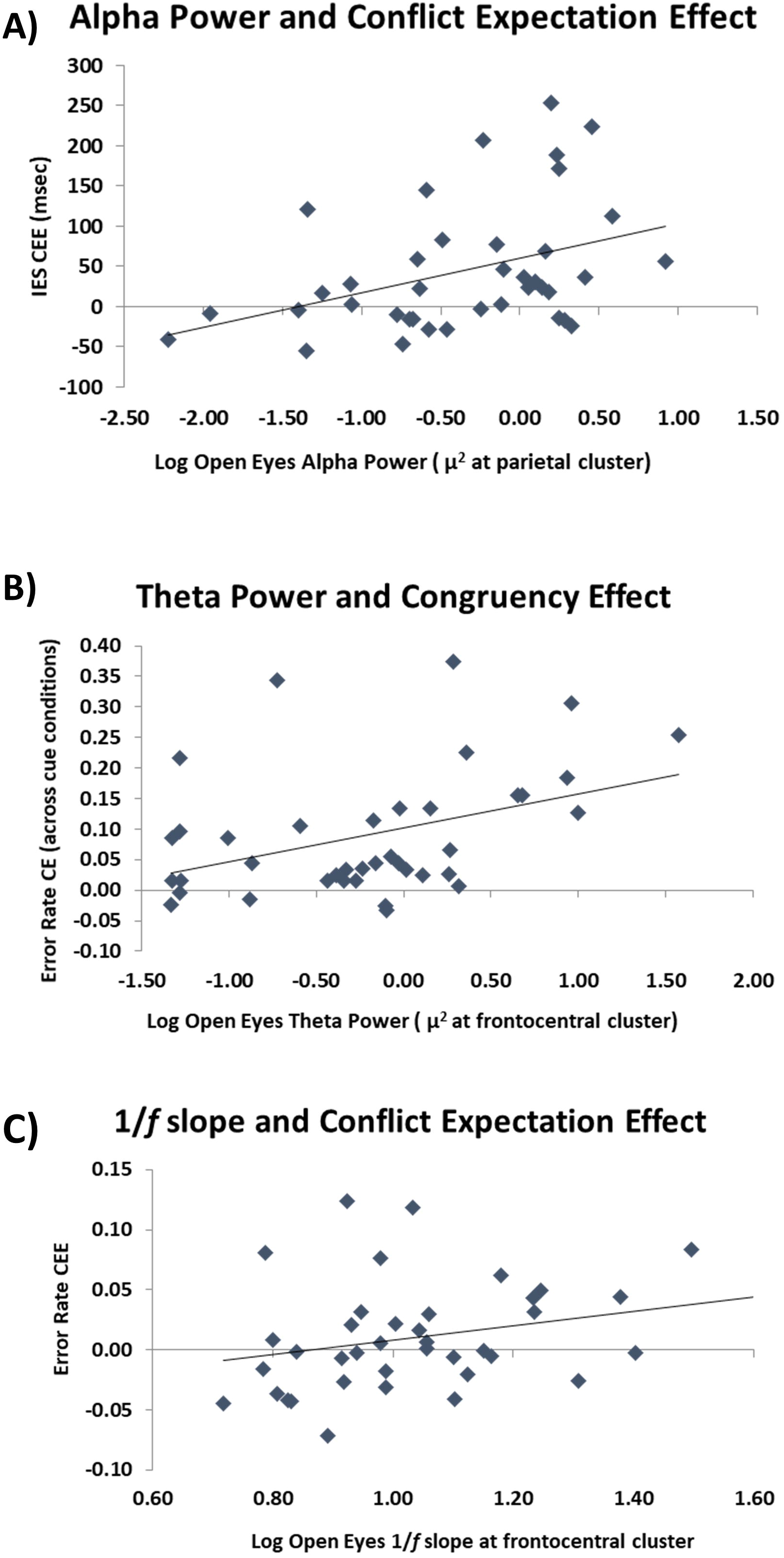
Relationship between EEG and cognitive control: **(A)** Scatterplot of parietal alpha power with the conflict expectation effect (CEE) on the inverse efficiency score (IES). The greater the alpha power, the more these individuals used the cue information (proactive control). **(B)** Scatterplot of frontocentral theta power with the congruency effect (CE) on error rate. The greater the theta power at rest, the greater the distractor interference (reactive control). **(C)** Scatterplot of frontocentral 1/*f* slope with the CEE on error rate. The steeper the 1/*f* slope, the more these individuals used the cue information (proactive control).

Exploratory analyses of 1*/f* slope indicated that it marginally predicted the error rate CEE, *r_s_*(37) = .271, *p* = .096, an effect which reached significance after controlling for age, *r_s_* (36) = .368, *p* = .023. This may indicate that higher levels of 1*/f* activity is predictive of greater proactive control (Figure 4C). Additionally, 1*/f* slope was not related to the error rate CE before, *r_s_* (37) = .233, *p* = .153, or after partialing out age, *r_s_* (36) = .119, *p* = .477. This suggests that 1*/f* slope may be more related to proactive rather than reactive control processes. It should be noted, however, that there was no clear *a priori* prediction for the 1*/f* slope, and that the effects reported were not corrected for multiple comparisons. Therefore, these findings require additional replications.

## 4. Discussion

Most (~90%) of the resting-state EEG power spectrum can be described by three parameters: the amplitude of the oscillatory spectral activity in the alpha and theta bands and the slope of the non-oscillatory 1*/f* component. These three spectral features were larger with closed than open eyes. Interestingly, two of these features differed between younger and older adults, such that older adults had less alpha power (at least with closed eyes) and less 1*/f* activity (i.e., less power at lower frequencies) than younger adults. Both of these findings are consistent with previous reports (Babiloni et al., 2006; Dave et al., 2018; Polich, 1997; Voytek et al., 2015). No age differences were found for theta activity, similarly to some previously reported research (Babiloni et al., 2006; Finely, Angus, van Reekum, Davidson & Schaefer, 2020). Consistent with our hypotheses, these electrophysiological characteristics were related to different aspects of cognitive control processing: increased alpha power (and, to a lesser extent, 1*/f* slope) predicted greater modulation of distractor processing given the cue information (conflict expectation effects), whereas theta power predicted greater distractor interference in error rates only (congruency effects). Following the dual mechanism framework proposed by Braver (2012), our evidence suggests that resting alpha power (and perhaps 1*/f* slope) are related to individual differences in proactive control processes, whereas theta power reflects individuals’ variations in reactive control. A compelling picture emerges from these data: separate parameters from the decomposed EEG power spectrum *at rest* may predict separable cognitive control processes during a behavioral task, independent of participant’s age. The alpha results resonate with previous work, which showed that alpha power predicted subsequent learning in a complex video game task involving multiple aspects of cognition (Mathewson et al., 2012).

Resting-state EEG has been previously used to investigate individual variability in cognitive status and cognitive performance, especially in the context of aging and cognitive decline. Most often, these recordings are conducted with eyes closed and have illustrated a shift in the EEG power spectrum as participants progress from a healthy cognitive status to MCI and AD (Babiloni et al., 2006; Kwak, 2006). This suggests that, as older adults change in cognitive status, their closed-eyes resting-state EEG spectrum changes concurrently, and that these changes can be used as indices of cognitive performance.

In the current study the predictability of the power spectrum characteristics on cognitive control processes occurred in the eyes open condition and in *both* younger and older adults, which suggests that this informative neural variability exists already in younger adults and may continue to provide predictive power as individuals age. This is not equivalent to saying that age has no effect on either alpha/theta amplitude or proactive/reactive control. Rather, it indicates that *mechanistic relationships* between the neural systems represented by alpha and theta and cognitive control functions may be maintained throughout the healthy lifespan, even though the individual component functions of these relationships may weaken (see Fabiani, 2012). As an example, in a study of the relationship between sequential effects in P300 amplitude and working memory function, we showed that the *relationship* was the same for younger and older adults, despite an evident reduction in working memory capacity and a shortening of the sequential activity in aging (Peltz, Gratton, & Fabiani, 2011).

As noted, there is an apparent dissociation between the typical age-related resting-state alpha differences observed with closed eyes, and the predictive role of alpha measures taken with open eyes, which is irrespective of age. This may indicate that individual differences in resting-state open-eyes alpha may better predict cognitive control phenomena occurring in tasks where visual attention is required, and eyes are open. It remains to be shown whether age-related differences in alpha obtained with closed eyes are instead more likely to be predictive of cognitive phenomena that do not rely on visual attention and can occur when the eyes are closed. This should be investigated in future research.

In many previous studies, alpha power fluctuations were investigated during ongoing cognitive control tasks using a time-frequency approach. These experiments often report alpha blocking during working memory encoding, attentional engagement, and proactive inhibition (Sauseng et al., 2005; Foxe & Snyder, 2011; Vissers et al., 2016; Wöstmann et al., 2019). They also report phasic posterior alpha power reductions following error trials (i.e., Cooper et al., 2016; van Driel, Ridderinkhof, & Cohen, 2012). These decreases in alpha power have been conceptualized as a mechanism associated with the refocusing of attention after an error occurred and when the updating of ongoing working memory task-related representations is needed. Gratton (2018; see also Mathewson et al., 2009; 2011) proposed that a temporary *blocking* of alpha is required for, or at least facilitates, the updating of representations. This is consistent with the idea that alpha per se is associated with the maintenance of existing representations over time (a phenomenon that would resist the updating process). This proposal links alpha mechanisms with the maintenance of representations, which would be required during proactive but not reactive control. This is consistent with the findings of the current study, in which alpha power at rest was predictive of the size of the conflict expectation effect (which requires maintenance of task-related representations in the interval between cues and response stimuli) but not of the congruency effect (which requires direct handling of conflict, without a delay).

According to Gratton (2018), bursts of frontocentral theta may provide a mechanism by which alpha is temporarily suspended to facilitate the updating of representations. As such, theta bursts would provide an opportunity for sustained alpha oscillations to be interrupted and for the maintenance of a representation to either be changed in favor of a new task-relevant representation, or in favor of a task-irrelevant distractor. Ancillary analyses, reported as **Supplementary Materials**, investigated whether theta occurs in a more burst-like manner than alpha (*at rest* and after accounting for 1/*f* activity). The results showed that this was indeed the case in the eyes open condition. These exploratory analyses suggest that, even during resting-state and in the absence of discernible external triggering events, theta tends to occur in shorter bursts than alpha. Further, the data suggest that the frequency of occurrence of theta bursts is strongly reflected by the total power across the whole resting-state recording period, whereas this is much less so for alpha. With respect to the functional significance of these bursts, we argue that, although a flexible engagement of theta interruption mechanisms is a useful process, integral to cognitive control, its excessive engagement may lead to maladaptive responses (i.e., distractibility) in conditions in which refocusing is not needed. This effect may be evident during resting-state situations: In such conditions, engagement of theta activity and blocking of alpha activity are not required, and the trait-like propensity to do so may be correlated with lower control abilities during tasks. It may also explain previously reported results, which indicate that IQ is negatively correlated with resting-state theta, and positively correlated with resting-state alpha (Doppelmayr et al., 2002; Jaušovec et al., 2001). Our results also inform the trait-like spectral differences in adults with ADHD, which indicate both increased theta power and decreased alpha power (Woltering, Jung, Liu, & Tannock, 2012), and could explain the tendency for increased distractibility in these individuals. This pattern may develop with the disorder, as Robertson et al. (2019) have shown that unmedicated children with ADHD have similar theta power but more alpha power compared to typically developing children. In the present study, such a propensity may be associated with increased susceptibility to the interference caused by incompatible flankers in the flanker task.

As mentioned previously, the evidence for age-related decreases in theta power is mixed (Babiloni et al., 2006; Cummins & Finnigan, 2007; Finely Angus, van Reekum, Davidson & Schaefer, 2020). Prichep et al. (2006) reported reductions in resting-state theta power in cognitively typical older individuals with subjective memory complaints seven years later, but only for those individuals whose mental status declined during the seven-year interval. This may address why younger and older adults in our study did not show theta power differences. Our older adults were cognitively intact and not reporting memory complaints. Finnigan and Robertson (2011) reported that relatively higher resting theta power in healthy older adults was related to better performance on verbal recall, attention, and executive function measures. Taken together these results suggest that the relationship between resting-state theta activity and cognitive abilities is complex, and perhaps not monotonic. Perhaps some minimum level of theta function needs to be maintained for appropriate cognition, but an excessive amount may signal a propensity to over-react to contextual changes and distraction, which may also be deleterious for cognition (e.g., Cools & D’Esposito, 2011). Further studies are needed to investigate this hypothesized inverse-U relationship between theta function at rest and cognition.

Although we did not find a difference in theta power with age, we did find a reduced 1*/f* slope with age, replicating similar age effects reported by Voytek et al. (2015) and Dave et al. (2018). It is important to note that, as computed in our study, 1*/f* slope is largely determined by low-frequency EEG activity (in a manner more similar to the intercept measures reported by Voytek et al., 2015). Thus, the reduction in 1*/f* slope observed in the current study fits well with other literature reporting reduced delta power across the adult lifespan (Babiloni et al., 2006; Polich, 1997). Interestingly, in both of these studies, alpha power was also shown to decrease with age, which we have reported as well. Here we show, for the first time, a relationship between 1*/f* slope and proactive control processes. This is of interest because it suggests that an additional electrophysiological mechanism may be involved in control processing. We have shown that alpha power and 1/*f* slope are not correlated with each other (in the open eyes condition), indicating that although they both predict proactive control processing, they are likely to be distinct electrophysiological signals. Since the analyses of 1/*f* were largely exploratory, they should be replicated and extended in future research.

Some limitations of the current study should be pointed out. First, our study is limited by its sample size. Nonetheless, predictive relationships were still present in this sample, suggesting that their effect size is sufficient. It is clear, however, that further testing with larger samples and other cognitive control tasks may be very useful to further validate the results reported here. Additionally, time-frequency analyses of EEG recorded during rest with different levels of alertness, and compared to diverse cognitive control tasks in a within-subject design would allow us to investigate the dynamics of these electrophysiological parameters and assess whether alpha and 1*/f* slope and theta power selectively mediate proactive and reactive control. These analyses may also be useful to demonstrate that EEG parameters measured at rest are predictive of possible trait-like individual differences in event-related time-frequency phenomena.

### 4.1 Conclusions

Resting-state EEG contains three dominant – and largely separable – electrophysiological signals: oscillatory alpha and theta power, and non-oscillatory 1*/f* slope. We found independent and separable predictive relationships between *resting-state* alpha power and proactive control, and theta power and reactive control, which existed regardless of participants’ age. The fact that these dimensions of cognitive control can be predicted from EEG activity *at rest,* and are therefore unrelated to specific task characteristics, suggests that they may represent important trait-like biomarkers. As such, they may prove useful in understanding life-span individual differences in cognition and may help researchers investigate variability in cognitive aging.

## Supporting information

Supplemental Theta Burst Analyses

## 5. Conflict of Interest

The authors declare that the research was conducted in the absence of any commercial or financial relationships that could be construed as a potential conflict of interest.

## 6. Author Contributions

GMC helped design the study, collected and analyzed the EEG data, and wrote the first draft of the manuscript; DCB collected and analyzed the behavioral data; MG helped with the EEG analysis pipeline and execution; KAL helped with experimental design and analyses; MF and GG designed the study, and supervised analyses and writing. All authors edited the manuscript and met to discuss results and theoretical implications.

## 7. Funding

This work was supported by NIA grant RF1AG062666 to G. Gratton and M. Fabiani.

## 8. Acknowledgments

We acknowledge Brooke Frazier, Dana Joulani, Rebecca Lii, Madeleine Peckus, Preeti Subramaniyan, and Yunsu Yu for help with data collection.

## Footnotes

1 The sample size reflects predicted effects for this task. To illustrate that the data is suitable for the analysis of individual differences we provide a reliability analysis on the EEG effects reported here.

2 Two older adults did not take the Shipley Vocabulary Scale, resulting in fewer degrees of freedom for this *t*-test.

3 To ensure that the correlations between even-odd epochs were not an effect of temporal continuity between power or 1/*f* in these epochs and to clarify whether power or 1/*f* changed as a result of participant fatigue, we also computed these correlations between the first and second half of the recording periods. They were all highly correlated: alpha power *r_s_*= .909, theta power, *r_s_*= .963; 1/f slope *r_s_*= .910. These correspond well to the even-odd reliability analysis reported in the results section.

## Notes

### Competing Interest Statement

The authors have declared no competing interest.

### Summary of Updates

Supplemental file added; author 3 added; additional analyses conducted

