## Supplemental Theta Burst Analyses for "Spontaneous alpha and theta oscillations are related to complementary aspects of cognitive control in younger and older adults"

### *Supplementary Material*

#### 1. Theta as a Burst: exploratory analyses

In the ancillary analyses reported here we focused again on resting-state conditions, to establish whether the timing patterns of theta and alpha occurrence are similar to those observed during tasks. During tasks, frontocentral theta is often reported to occur in short-lived bursts. Specifically, theta bursts index brief (< 500 ms) brain oscillatory responses that occur after the onset of attention-catching stimuli (Cavanagh & Frank, 2014; Cohen & Donner, 2013). Alpha, instead, is often present before stimulus onset, with theta bursts typically occurring simultaneously with the alpha suppression that follows stimuli that capture attention.

Here, we sought to quantify the presence or absence of burst-like theta activity *at rest,* and compared it to alpha activity, which we assumed to occur in a more sustained manner compared to theta. We segmented the eyes-open recording into 1024 ms epochs to increase temporal resolution, computed a fast Fourier transform and detrended the 1/*f* slope from each spectrum using the methods described in this paper. An FFT was used (rather than an FFT with Welch's method) to decompose the spectra, in order to avoid potential distortions of the effects introduced by the Hamming window incorporated in Welch's method.

If resting-state theta occurs in bursts, it should be present only a small amount of the time, or for very few contiguous epochs. If resting-state alpha is more sustained compared to theta, it should be present in the majority of the epochs. However, some of the power observed in the theta (or alpha) band may be determined by the contribution of the 1/*f* slope within that band (estimated using the same logic as in the rest of the paper). Therefore, the "true" power of theta (or alpha) for each epoch is given by the difference in power between the raw power for the theta (or alpha) band and the power for that band predicted by 1/*f*. If this difference is positive, it would indicate that theta (or alpha) is present for that specific epoch; if the difference is 0 or negative, it would indicate that theta (or alpha) did not occur in that epoch.

Using this approach, under the null hypothesis (power of theta is equal to 0), the base rate of epochs showing theta (or alpha) activity (after accounting for 1/*f)* should be 50% (P(null) = .50). Therefore, for each individual we can estimate the corrected proportion of points with signal (theta or alpha oscillations) present as:

$$P(true\ signal) = \frac{P(signal > 1/f) - P(null)}{1 - P(null)}$$

We found that the corrected probability that a particular epoch shows theta (averaged across individuals) is 24.0% $\pm$ 6.5% with a 95% CI [10.8%, 37.2%]. In contrast, the corrected probability that a particular epoch shows alpha is 63.8% $\pm$ 6.7% with a 95% CI [50.2%, 77.5%]. The comparison between the corrected probabilities for theta and alpha power was significant, $t(38) = 5.13$, $p < .00001$. These data suggest that theta occurs on average during approximately 25% of the epochs whereas alpha occurs significantly more often (approximately 64%). This appears to support the idea that theta at rest occurs in bursts, even in the absence of discernible external triggering events. Instead. alpha was more continuously present.

For neither alpha nor theta the frequency of epochs with power greater than what was predicted by the null hypothesis was correlated with age ($r$ = .036 for alpha and $r$ = .054 for theta). Finally, we also assessed the relationship (across individuals) between alpha and theta average power across the entire period of recording and the frequency of epochs with power greater than what was predicted by the null hypothesis. A strongly positive correlation ($r$ = .566, $p$ = .0002) emerged for theta, but the correlation was much smaller for alpha ($r$ = .335, $p$ = .040). This finding is consistent with the idea that the average theta power during the resting state period is much more dependent on the frequency of occurrence of short-duration activity (“bursts”) than alpha power, as it could predicted on the basis of Gratton’s (2018) model.
